# Mitochondrial depolarization stabilizes the vitamin B12 chaperone MMADHC in the cytosol to increase MTR activity

**DOI:** 10.64898/2025.12.31.697091

**Authors:** Sneha P. Rath, Zhu Li, Arkajit Guha, Fangcong Dong, Ruma Banerjee, Vamsi K. Mootha

## Abstract

Of the ∼1100 mitochondrial proteins, only a handful like PINK1 and ATFS-1 are known to stabilize and relocalize upon collapse of the proton motive force (PMF) to execute signaling roles. To systematically identify genes that increase exclusively at the protein level upon PMF collapse, we performed a joint proteomic and RNA-seq screen. The screen revealed 10 candidates (six mitochondrial), including the vitamin B12 chaperone MMADHC and cytosolic B12-dependent methionine synthase (MTR). MMADHC is short-lived across cell types and we show that its levels increase with PMF collapse. MMADHC stabilization precedes PINK1 activation in a time course of increasing mtDNA depletion, suggesting greater sensitivity to PMF collapse. MMADHC accumulates in mitochondria with LONP1 inhibition but in the cytosol upon PMF collapse, likely due to mitochondrial import failure. Cytosol-stabilized MMADHC increases MTR levels and activity. Altogether, the mitochondrial PMF regulates the cytosolic B12-dependent MTR, integral to one-carbon metabolism, by controlling the stability and compartmentalization of the B12 chaperone MMADHC.

**Significance Statement:** Humans have only two vitamin B12-dependent enzymes – mitochondrial MMUT and cytosolic MTR – and both require a common B12 chaperone MMADHC. We discover that MMADHC is a low abundant, short-lived protein that is continuously imported and degraded by energized mitochondria. Upon collapse of the mitochondrial proton motive force, MMADHC accumulates in the cytosol and increases the levels and activity of MTR, critical for one-carbon metabolism. This PMF-dependent regulation of MMADHC stability and localization is important for understanding cofactor rationing and spatiotemporal compartmentalization of B12 metabolism.

## Introduction

The mitochondrial proteome comprises ∼1100 nuclear-encoded proteins^1^ that are translated on cytosolic ribosomes and imported into mitochondria. Depending on the final sub-mitochondrial destination, protein import requires chaperones, redox recycling, proton motive force (PMF), and ATP^2^. Thus, the mitochondrial proteome, which represents ∼5% of the total human proteome, is poised to sense mitochondrial proteostasis, redox, and bioenergetic status simply by virtue of its dependence on import.

Three prominent examples where cells exploit the opportunity of import-dependent mitochondrial surveillance and couple it to signaling are PINK1, *C. elegans* ATFS-1, and DELE1. PINK1 and ATFS-1 constantly undergo PMF-dependent degradation: PINK1 is clipped by PARL^3^ during its transit into the mitochondrion and released to the cytosol for proteasomal degradation, whereas ATFS-1 is degraded by LONP1^4^ in the mitochondrial matrix. At a critical threshold of PMF collapse, both proteins fail to import, and while PINK1 accumulates on the OMM to initiate mitophagy^5^, ATFS-1 translocates to the nucleus to activate the mitochondrial unfolded protein response^6^. More recently, DELE1 was also shown to sense PMF collapse, as well as other stresses, during transit through the OMM and IMM, which causes its mitochondrial release and subsequent activation of the cytosolic integrated stress response kinase HRI^7,8,9^. An open question is whether other mitochondrial proteins exhibit similar PMF-dependent regulation and the impact of their conditional extra-mitochondrial sequestration or retro-translocation.

Here, we performed a multi-omic screen to systematically identify mitochondrial proteins that are stabilized upon PMF collapse. Our analysis spotlights MMADHC, a vitamin B12 chaperone. Humans have two B12-dependent enzymes: (i) cytosolic methionine synthase (MTR) and (ii) mitochondrial methylmalonyl-CoA mutase (MMUT)^10^. MTR converts methyl-tretrahydrofolate, the circulating form of folate, to tetrahydrofolate (THF) which is required for nucleotide biosynthesis, and methylates homocysteine to produce methionine in the process^11^. MMUT converts methylmalonyl-CoA to succinyl-CoA, catalyzing the final step in the TCA anaplerosis of valine, methionine, isoleucine, threonine and odd-chain fatty acids^12^. MMADHC is genetically linked to both MTR and MMUT based on patient mutations that lead to a toxic accumulation of their respective substrates, causing either isolated or combined homocystinuria and methylmalonic aciduria^13^. Human MMADHC has an N-terminal mitochondrial targeting sequence and is a 296 amino acid long protein, the first third of which is predicted to be disordered while the last two-thirds belongs to the ferredoxin nitroreductase superfamily and coordinates B12^14^. While the role of MMADHC in the mitochondrial B12 pathway remains elusive, previous studies have demonstrated that it can load the B12 cofactor onto the cytoplasmic target MTR^15^. Whether and how the mitochondrial versus cytoplasmic compartmentalization of MMADHC is regulated remains largely unexplored.

Using an unbiased approach, our study uncovered that MMADHC is regulated by the mitochondrial PMF. We show that MMADHC is continuously degraded by LONP1 in energized mitochondria but upon PMF-collapse it is retained in the cytosol where it is responsible for increasing MTR levels and activity. Thus, the PMF regulates MMADHC stability and compartmentalization and, in turn, cytosolic MTR activity, with implications for one-carbon metabolism.

## Results

### A multi-omic screen identifies MMADHC as the top hit that increases in response to mitochondrial PMF collapse

To systematically identify genes that are upregulated only at the protein level in response to a collapse of the mitochondrial PMF, we performed TMT proteomics and RNAseq analysis of K562 cells treated with antimycin and oligomycin (“anti+oligo”) for 3 days. Of the 6378 genes with combined protein and RNA level data, only 10 genes were up-regulated exclusively at the protein level (cutoff >2-fold). Six of these are mitochondrial proteins, including CHCHD2 which is previously reported to increase at the protein level upon mitochondrial depolarization^16^ (figure 1A). In contrast, ATF4 target genes were induced both at the RNA and protein level, as expected for a transcriptional response. Although PINK1 was not detected, we observed a robust increase in the PINK1-dependent phosphosite pSer65-ubiquitin^17^ in response to anti+oligo (figure 1B). We focused on the top hit that increased only at the protein level, the vitamin B12 chaperone MMADHC, and note that the cytosolic B12-dependent MTR, which interacts physically with MMADHC^13^, is also on this list.

**Figure 1:**
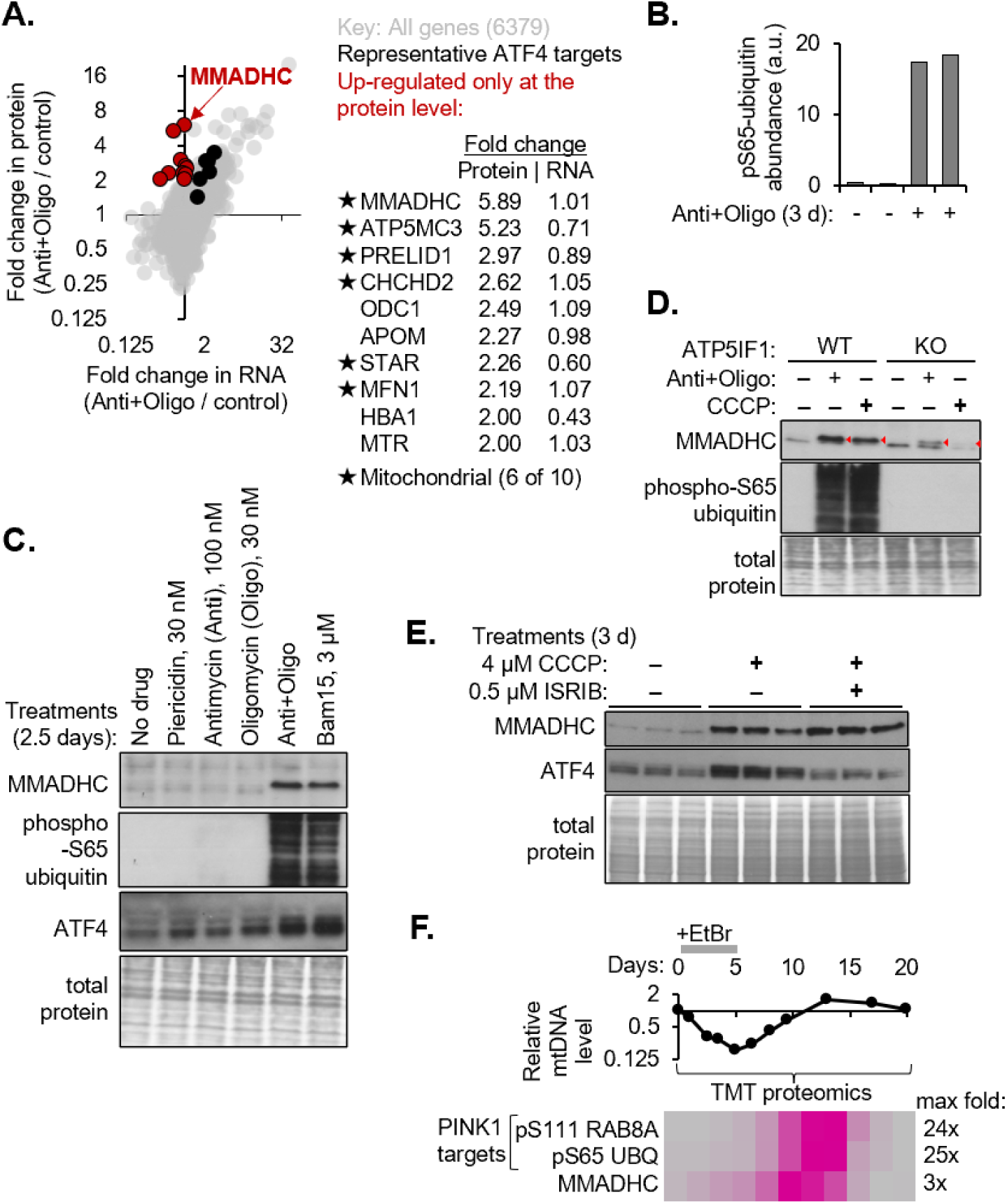
A multi-omic screen identifies MMADHC, a short-lived protein that increases with collapse of the mitochondrial proton motive force (A) Scatter plot of average fold change in protein vs RNA levels measured by TMT proteomics and RNAseq respectively in K562 cells treated with antimycin and oligomycin (anti+oligo) for 3 days, in two biological replicates. Black points show representative ATF4 target genes ASNS, CTH, PHGDH, PSAT1, PSPH, TRIB3. Red points are genes with fold change in protein >2 and RNA <1.2-fold and are listed in the table adjacent to the scatter plot. (B) Abundance of the PINK1-dependent phosphosite pSer65-ubiquitin detected by TMT proteomics in the samples in A (C) Western blot analysis of the indicated target genes in K562 cells treated with electron transport chain inhibitors piericidin, antimycin, or oligomycin, targeting complexes I, III, and V respectively, or drugs that collapse the proton motive force, ‘anti+oligo’ or the protonophore BAM15. Blot is representative of three biological replicates. (D) Western blot analysis of MMADHC and phospho-S65 ubiquitin in sgGFP or sgATP5IF1 K562 cells treated with antimycin and oligomycin (anti+oligo) or CCCP for 3 days. Total protein stained with Coomassie for loading control. Blot is representative of two biological replicates. (E) Western blot analysis of MMADHC and ATF4 in K562 cells treated with uncoupler CCCP or co-treated with CCCP and ISRIB in three biological replicates. Total protein stained with Coomassie for loading control. (F) QPCR for mtDNA normalized to nuclear DNA in K562 cells treated with ethidium bromide (EtBr) for 5 days, followed by EtBr-withdrawal for 15 days. Row-scaled heatmap of protein levels of MMADHC vs two PINK1 target phosphosites quantified by TMT proteomics at the same timepoints as mtDNA analysis.

To distinguish whether MMADHC is upregulated in response to electron transport chain (ETC) inhibition or mitochondrial PMF collapse, we compared MMADHC levels across cells treated with a variety of mitochondrial poisons. MMADHC levels did not increase in response to inhibitors of ETC complexes I, III, or V (piericidin, antimycin A, and oligomycin respectively, figure 1C). MMADHC, but not its mRNA, was up-regulated in response to anti+oligo as well as the mitochondria-specific protonophore BAM15 (figure 1C). Both these treatments collapsed the PMF as confirmed by accumulation of a PINK1 target, phospho-S65 ubiquitin (figure 1C). In contrast to MMADHC, ATF4 was induced to varying degrees in all treatments, as expected^18,19^ (figure 1C).

To further confirm that the increase in MMADHC is responsive to PMF collapse, we tested the effect of ATP5IF1 knockout (KO), which defends the PMF via complex V reversal when the membrane depolarization exceeds a certain threshold^20^. Control cells treated with anti+oligo or CCCP for three days showed the expected increase in MMADHC and the PINK1 target phospho-S65 ubiquitin, both of which were strongly blunted or prevented altogether in ATP5IF1 KO cells (figure 1D).

The increase in MMADHC exclusively at the protein level might be due to enhanced translation or decreased proteolysis – mechanisms which are known to regulate ATF4 and PINK1 respectively. In ATF4, ribosomes engage at the conserved upstream open reading frames (uORFs), while the downstream ORF is translated only upon eIF2α phosphorylation, which limits translation initiation and promotes ribosome scanning past the uORFs^21^. In principle, human MMADHC could be a candidate for such regulation because it has an uORF which is conserved in *M. musculus*, *X. laevis*, *D. melanogaster*, as well as the sea slug *A. californica* (figure S1). However, ISRIB treatment to rescue translation despite eIF2α phosphorylation^22^ blunted induction of ATF4, but not that of MMADHC, in response to the mitochondrial uncoupler CCCP (figure 1E), ruling out translational upregulation as a mechanism for MMADHC regulation.

Next, we compared the kinetics of MMADHC increase relative to PINK1-dependent phosphosites. Proteomics analysis of K562 cells during a time course of EtBr treatment^23^, which depletes mtDNA, followed by EtBr withdrawal to recover mtDNA copy number, revealed that the onset and peak of increase in MMADHC preceded that of the PINK1 targets, phosphosites pS111 on RAB8A and pS65 on UBQ (figure 1F, dataset is part of a separate submitted manuscript). These data reveal that MMADHC is more sensitive to PMF collapse than PINK1.

### MMADHC is among the shortest-lived proteins in multiple cell types

Analysis of previously published proteome-wide half-lives^24^ from cycloheximide pulse and proteomics experiments indicates that MMADHC, with a half-life of ≤1 h, is among the top 50 shortest-lived proteins across all four cell lines/lineages tested (figure 2A). Notably, three other hits from our screen (figure 1A) are also short-lived in at least one cell line: PRELID1 (<1 h), ODC1 (<1 h), and CHCHD2 (∼2 h). Our joint RNA-seq and TMT proteomics of K562 cells revealed that, although MMADHC mRNA is highly expressed, the steady-state levels of the protein are low. Thus, while *MMADHC* mRNA ranks 258^th^, the protein ranks 4445^th^ out of 6378 genes ranked in descending order of RNA or protein expression respectively (figure S2A, S2B). There is a strong correlation between mRNA and protein levels across the 6378 genes with joint data (Spearman correlation: 0.38, p value: 1.1E-222). For example, subunits of the cytosolic ribosome are highly expressed at both the RNA and the protein level in contrast to MMADHC (figure S2C).

**Figure 2:**
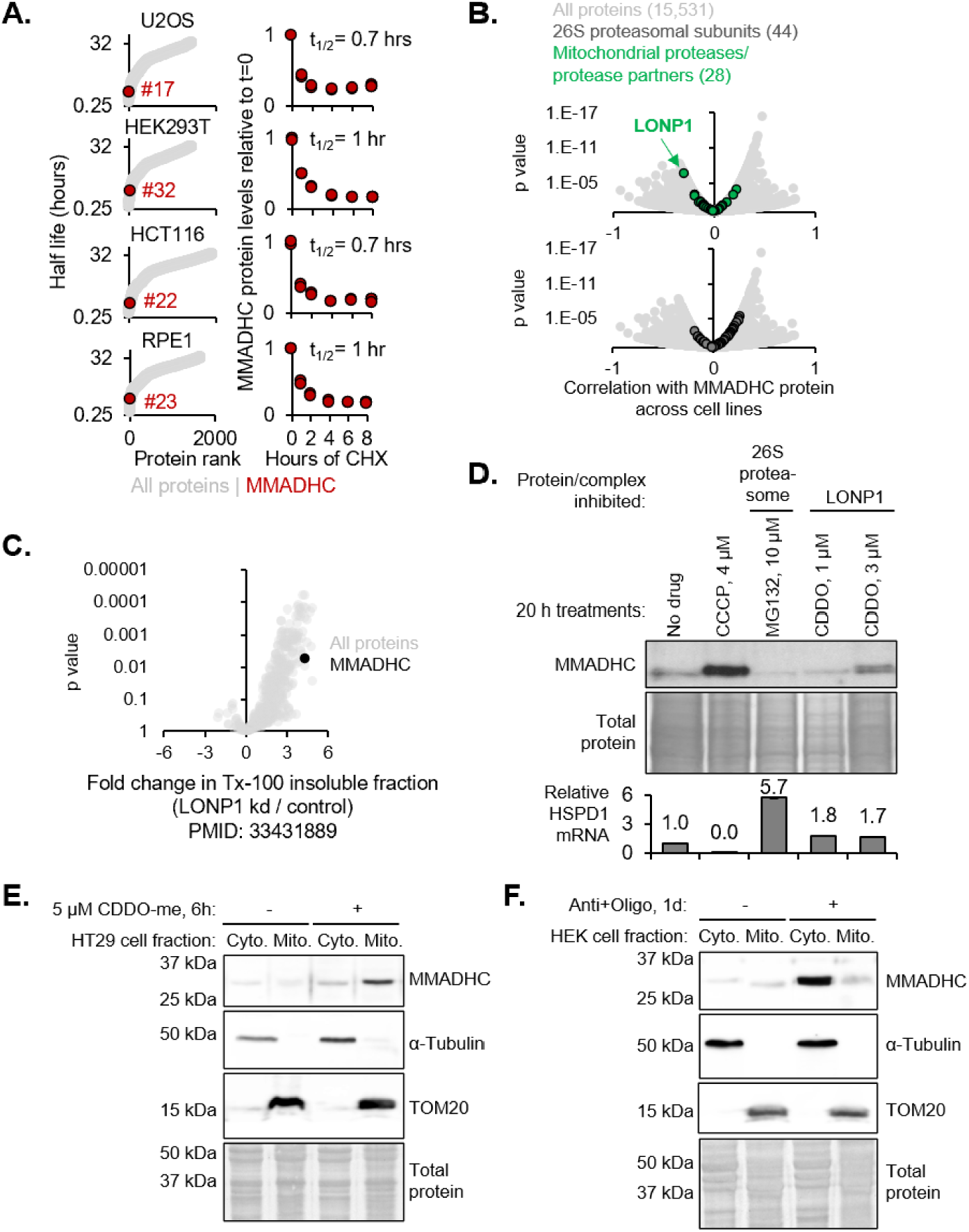
**MMADHC accumulates in mitochondria with lonp1 inhibition and in the cytosol with mitochondrial depolarization** (A) Protein half-lives (left) and MMADHC levels (right) measured by cycloheximide (CHX) chase in four cell lines. Re-analysis of data from PMID: 34626566. (B) Volcano plots of MMADHC protein correlated to all other proteins across CCLE. Re-analysis of TMT proteomics data from PMID: 31978347. (C) Volcano plot of fold change in protein levels in the triton-100 insoluble fraction in response to LONP1 knockdown. Re-analysis of raw data from PMID 33431889. (D) Western blot of MMADHC in K562 cells treated with CCCP, proteasomal inhibitor MG132, and LONP1 inhibitor CDDO-me. Total protein is stained with Coomassie for loading control. QPCR quantification of HSPD1 mRNA relative to ACTB as a proxy of proteotoxic stress. (E) Western blot of cytosolic and mitochondrial fractions of HT29 cells treated with LONP1 inhibitor CDDO-me. Total protein is stained with Ponceau for loading control. Blot is representative of three biological replicates. (F) Western blot analysis of cytosolic and mitochondrial fractions of HEK293 cells treated with antimycin and oligomycin for one day. Blot is representative of three biological replicates.

### MMADHC accumulates in mitochondria upon LONP1 inhibition but in the cytosol upon mitochondrial depolarization

We hypothesized that MMADHC might be regulated like PINK1 or ATFS-1, which are constantly imported and degraded by PARL/proteasome or LONP1 respectively and stabilized only when they fail to import due to mitochondrial depolarization. To test this hypothesis, we first sought to identify the protease responsible for MMADHC turnover.

We performed correlation analysis of MMADHC against each of ∼15,000 detected proteins across the Cancer Cell Line Encyclopedia (CCLE)^25^. Out of the 28 detected mitochondrial proteases, LONP1 is the sole protease that is significantly negatively correlated with MMADHC while in contrast, the cytosolic proteasomal subunits do not show any significant correlation with MMADHC (figure 2B). Breaking down the CCLE analysis by lineage, MMADHC is negatively correlated with LONP1 within several lineages (figure S3A), most notably in the ovarian and colon lineages (figure S3B).

Re-analysis of published proteomics^26^ of the triton-100 insoluble fraction from control versus LONP1 knockdown cells revealed that MMADHC was significantly enriched in the insoluble fraction upon LONP1 knockdown (figure 2C), consistent with a putative role of LONP1 in degrading MMADHC and preventing its aggregation in mitochondria. Chemical inhibition of LONP1 by CDDO-me increased MMADHC in K562 cells, but proteasomal inhibition by MG132 did not (figure 2D). Our data is consistent with the prior observation that LONP1 inhibition increased MMADHC in H1944 cells^27^.

Next, we determined MMADHC localization in response to LONP1 inhibition versus mitochondrial depolarization. Because CDDO-me treatment decreased K562 cell viability, we tested its effect in a different cell line, HT-29, selecting from the colon lineage where LONP1 is strongly and negatively correlated with MMADHC (figure S3B). In HT-29 cells treated with CDDO-me, MMADHC increased and was enriched in the mitochondrial fraction (figure 2E), consistent with MMADHC being degraded primarily by LONP1 in the mitochondrial matrix. In contrast, MMADHC accumulated in the cytosol upon mitochondrial depolarization by anti+oligo (figure 2F). Thus, MMADHC is regulated in a PINK1-like manner in that its turnover depends on a mitochondrial protease and PMF and it is stabilized in the cytosol following PMF collapse.

### Cytoplasmic stabilization of MMADHC increases MTR protein and activity

To determine the role of MMADHC stabilized by depolarization, we performed TMT proteomics on control and MMADHC KO cells treated with or without anti+oligo. Analysis of 7,376 total proteins revealed only one protein aside from MMADHC that increased with anti+oligo in control cells but failed to do so in MMADHC Kos, i.e. MTR (figure 3A). No other proteins were differentially elevated or depleted by anti+oligo in control versus MMADHC KO (figure 3A). This data allows us to draw a causal genetic link between the increase in MTR and MMADHC by mitochondrial depolarization, both of which were also hits in our initial screen (figure 1A) and conclude that the increase in MTR upon PMF collapse is MMADHC-dependent. Conversely, however, the increase in MMADHC upon PMF collapse is MTR-independent i.e. MMADHC increases comparably in response to anti+oligo in control and MTR KO cells (figure S4A). Furthermore, MTR is attenuated in MMADHC KO even in the absence of anti+oligo (figure 3B), indicating that MMADHC influences basal levels of MTR.

**Figure 3.**
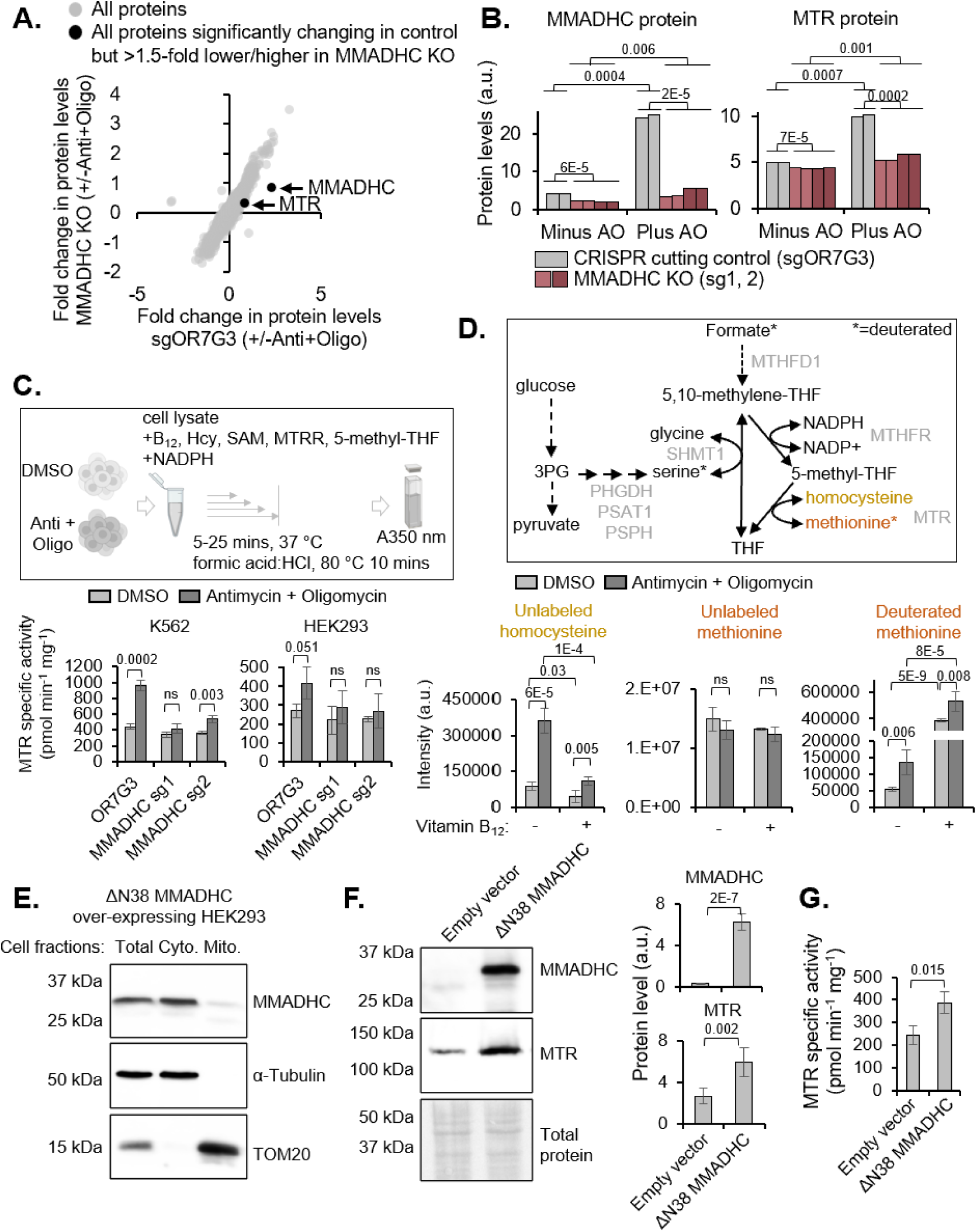
**Cytosol-stabilized MMADHC increases MTR protein and activity** (A) Scatter plot of fold change in protein levels of MMADHC KO (sg1, sg2 combined) vs control (sgOR7G3 CRISPR cutting control) cells treated with antimycin and oligomycin (anti+oligo) for three days and analyzed by TMT proteomics. Average fold from two replicates for control (sgOR7G3) and four replicates across two separate sgRNAs for MMADHC are plotted. (B) Bar plots of MMADHC and MTR levels from the experiment shown in (A) (C) Schematic of MTR activity assay in cell extracts. MTR specific activity measured in extracts of K562 and HEK293 cells treated with or without antimycin and oligomycin for three days. Error bars represent standard deviation across three biological replicates. (D) Schematic of deuterated formate tracing. Quantification of MTR substrate homocysteine and product methionine (unlabeled and deuterated) by mass spectrometry in K562 cells treated with or without antimycin and oligomycin for three days and labelled with fully deuterated formate in the presence or absence of vitamin B12 for 9 hours. Error bars represent standard deviation across four biological replicates. (E) Western blot analysis of MMADHC in total lysate, cytosolic, and mitochondrial fractions of HEK293 cells transiently over-expressing MMADHC lacking the first 38 a.a. (MMADHCΔN38). Blot is representative of three biological replicates. (F) Western blot analysis of MMADHC and MTR in HEK293 cells stably over-expressing empty vector or MMADHCΔN38. Total protein is stained with Ponceau for loading control. Bar plots on the right show western blot quantification, error bars represent standard deviation across five biological replicates. (G) MTR specific activity measured as in ‘C’ in HEK293 cells transiently over-expressing empty vector or MMADHCΔN38. Error bars represent standard deviation across three biological replicates.

Next, we examined the functional impact of mitochondrial depolarization-driven and MMADHC-dependent increase in MTR. We determined MTR activity in cell lysates in the presence of the MTR substrates, homocysteine and 5-methyl-tetrahydrofolate (THF), and a reactivation system (MTRR, NADPH, S-adenosylmethionine) (figure 3C, schematic). We observed a 1.5-2-fold increase in MTR specific activity in lysates of anti+oligo-treated cells, which was ablated by MMADHC KO in cells from two lineages, K562 & HEK293 (figure 3C). Western blot analysis of the same samples confirmed the MMADHC-dependent increase in MTR upon mitochondrial depolarization in both cell lines (figure S4B, S4C).

Additionally, we assayed MTR activity using deuterated formate to quantitatively trace the conversion of 5,10-methylene-THF to THF (figure 3D, schematic). In this experiment, we quantified: (i) methionine labeling via the activities of MTHFR and MTR and (ii) serine labeling via the activities of SHMT1/2 in cells cultured in the presence of homocysteine to ensure it was not limiting for MTR activity. In addition, cells were also cultured in the absence or presence of exogenous vitamin B12 to distinguish holo-MTR (that was already loaded with vitamin B12 in cells) from total MTR.

The deuterated formate tracing experiment revealed that homocysteine accumulates with anti+oligo treatment and is largely rescued by B12 supplementation (figure 3D). The presence of excess B12 in the culture medium is expected to support B12 loading independently of MMADHC, thus enabling assessment of total MTR activity. Concomitantly, deuterated methionine, which is indicative of MTR activity, increased in response to anti+oligo, independent of B12 supplementation (figure 3D) while the levels of unlabeled methionine were unchanged. B12 supplementation boosted deuterated methionine levels ∼10-fold even without anti+oligo, while a smaller three-fold increase was observed in the presence of anti+oligo (figure 3D). Finally, deuterated formate tracing to serine, depends on the conversion of 5,10-methylene-THF to THF via SHMT1/2and does not require B12. Anti+oligo increased unlabeled serine but decreased deuterated serine levels (figure S5), suggesting SHMT1/2 reversal^16,28^. Collectively, the *in vitro* MTR activity and deuterated formate tracing reveal an increase in MTR activity upon mitochondrial depolarization and highlights that MTR exists largely in the apo form, limited by B12 loading.

To investigate if cytosolic stabilization of MMADHC in the absence of mitochondrial depolarization also increases MTR levels and activity, we characterized a ΔN38 variant of MMADHC that lacks the mitochondrial targeting sequence (MTS). The longest predicted MTS for human MMADHC across TargetPv1.0^29^, v2.0, TPpred2^30^, and MitoFates^31^ is 38 residues. Unlike the native protein, ΔN38-MMADHC was readily detected by western blot analysis even in the absence of anti+oligo and was localized almost exclusively in the cytosol (figure 3E). Over-expression of ΔN38-MMADHC increased MTR levels (figure 3F) and activity (figure 3G), demonstrating that cytosolic MMADHC is sufficient to boost both.

## Discussion

In this study, we sought to systematically discover proteins that, like PINK1 or ATFS-1, are stabilized upon mitochondrial PMF collapse. Our joint proteomic and RNA-seq screen in K562 cells identified 10 genes (six mitochondrial) that are up-regulated exclusively at the protein level in response to PMF collapse. We followed up on the top hit, the vitamin B12 chaperone MMADHC, and demonstrated that its short half-life is driven by its continual import into energized mitochondria and degradation by LONP1. We summarize our model for MMADHC regulation in Figure 4. MMADHC can be conditionally stabilized in two cellular “zip-codes”: in mitochondria with LONP1 inhibition or in the cytosol with PMF collapse (figure 4A), where it increases the levels and activity of the cytosolic B12-dependent MTR (figure 4B). An open question is whether this stabilization and boost in cytosolic MTR activity is relevant in any specific physiological or disease state.

**Figure 4:**
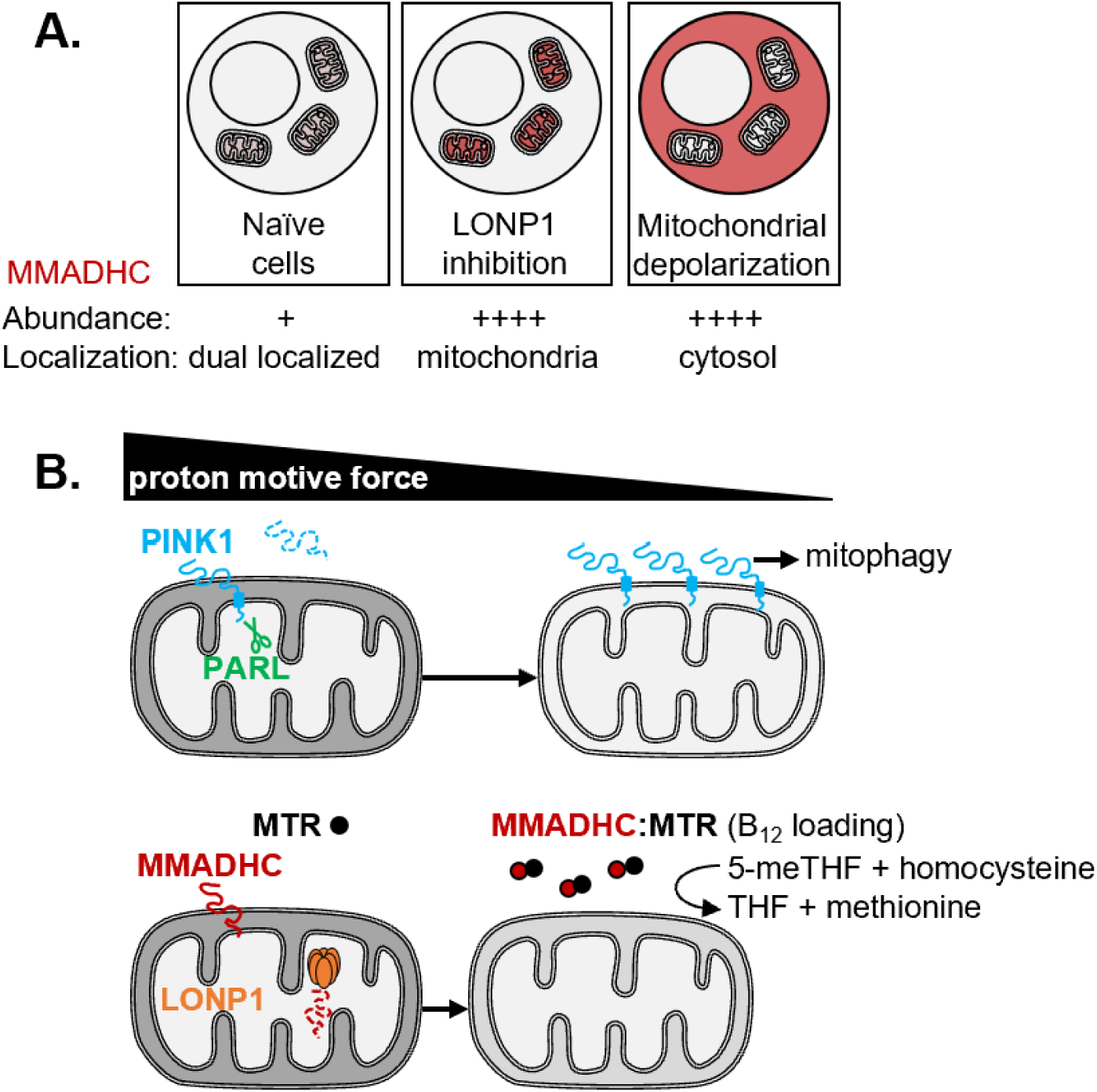
**Model for the regulation of MMADHC by the mitochondrial proton motive force and LONP1** (A) Context-dependent MMADHC abundance and localization: MMADHC is present at very low steady state levels in cells with healthy mitochondria but accumulates in mitochondria with LONP1 inhibition and in the cytosol with mitochondrial depolarization. (B) MMADHC is stabilized earlier than PINK1 upon PMF collapse and accumulates in the cytosol where it boosts the levels and activity of the B12-dependent MTR.

The two human vitamin B12-dependent enzymes reside in separate compartments (MMUT in mitochondria, MTR in the cytosol) with MMADHC being the only protein common to both branches after B12 is exported from the lysosome. Thus, regulating the distribution and stability of MMADHC likely allows cells to ration cofactor availability for each branch and to spatiotemporally compartmentalize vitamin B12 metabolism. In murine hematopoetic stem cells (HSCs), mitochondrial turnover of Mmadhc by Afg3l2, the highest-expressed mitochondrial protease in HSCs, restricts B12-dependent TCA cycle anaplerosis and maintains HSC homeostasis^32^. Further investigation is needed to systematically determine how different cell types/biological contexts influence MMADHC stability and localization to regulate B12 metabolism. For example, highly proliferative cells that rely on MTR to acquire the THF needed for nucleotide synthesis^33^ may have higher levels of cytosolic MMADHC, while tissues such as liver, which has the highest expression of MMUT (GTEx Analysis Release V10, dbGaP Accession phs000424.v10.p2), may achieve relatively higher levels of mitochondrial MMADHC by favoring its import or potentially slowing its degradation.

The continuous mitochondrial turnover and conditional cytosolic retention of MMADHC upon PMF collapse allows cells to couple increased MTR activity to PMF collapse. We note that the immediate metabolic ‘neighborhood’ of MTR includes pathways already known to be transcriptionally elevated by ETC dysfunction via the ATF4 response, namely serine synthesis, transsulfuration, and xCT. We now report a distinct, ATF4-independent, and post-transcriptional mechanism of increasing MMADHC, and consequently MTR, specifically upon PMF collapse but not with other ETC inhibition. We speculate that increased MTR is needed upon PMF collapse because of defective homocysteine clearance and SHMT1/2 reversal revealed by our formate tracing experiment. The latter is consistent with our previous study which identified serine as the top increasing metabolite in mtDNA-depleted cells and SHMT2 reversal upon ETC inhibition^16^.

Cells exploit the conditional stabilization and localization of mitochondrial proteins not only for signaling stress, as in the case of PINK1, ATFS-1, and DELE1, but also for modulating metabolism. Besides MMADHC, our screen also recovered STAR, the steroidogenic acute regulatory protein, which functions at the OMM to facilitate the rate-limiting import of cholesterol to the IMM^34^. Like MMADHC, STAR is also rapidly degraded by LONP1^35^ in the mitochondrial matrix and this turnover is thought to be crucial for regulating steroid synthesis. In uricotelic (uric acid-excreting) vertebrates like chickens, another metabolic enzyme is differentially mito-localized: glutamine synthetase (GS) is cytosolic in astrocytes but mitochondrial in hepatocytes where it is critical for detoxifying ammonia. This differential localization is achieved by GS harboring a weak MTS and hepatocyte mitochondria specifically having a high enough PMF to drive GS import^36^.

The importability of mitochondrial proteins, which is dictated by intrinsic protein features such as MTS strength and the bioenergetic state, appears to be finely tuned for each protein’s function and organelle homeostasis. We observed that PINK1 is more resilient to PMF collapse than MMADHC likely due to PINK1 having the stronger MTS, which allows cells to reserve PINK1-dependent mitophagy only for severe mitochondrial depolarization. A comparative study of MTS strengths in yeast uncovered remarkable variability in the import kinetics of proteins not known to be involved in stress signaling^37^. We therefore speculate many additional mitochondrial proteins than currently appreciated to be stabilized/(re)localized at specific PMF thresholds.

## Materials and Methods

### Cell culture

K562 (female), HEK293 (female), and HT29 (female) cells were obtained from the ATCC and cultured in DMEM (Gibco 11995-065) with 10% FBS (Sigma Aldrich F2442) and 100 U/mL penicillin/ streptomycin at 5% CO2 and 37°C. For experiments with electron transport chain inhibitors and uncouplers, cells were supplemented with 50 μg/mL uridine. When necessary, K562 cells were selected with 2 μg/ml puromycin (GIBCO). Cell lines were authenticated by STR profiling (ATCC) and tested to ensure the absence of mycoplasma once every 3 months. For experiments with CRISPR-Cas9 knockout, polyclonal cells were used throughout.

### TMT proteomics

Quantitative proteomics was performed at the Thermo Fisher Scientific Center for Multiplexed Proteomics (Harvard). **Sample preparation for mass spectrometry:** Cell pellets were lysed in 8M urea, 200 mM EPPS pH 8.5, protease and phosphatase inhibitors. 150 ug of protein from each sample was reduced with TCEP, alkylated with iodoacetamide, and then further reduced with DTT. Proteins were precipitated onto SP3 beads to facilitate a buffer exchange into digestion buffer. Samples were digested with Lys-C (1:50) overnight at room temperature and trypsin (1:50) for 6 hours at 37°C. Peptides were labelled with TMTPro reagents. 2 uL of each sample was pooled and used to shoot a ratio check in order to confirm complete TMT labelling. All 18 TMTPro-labelled samples from each group were pooled and desalted by Sep-pak. Phosphopeptides were enriched using a Peirce High-Select Fe-NTA Phosphopeptide Enrichment Kit. The flow through from the enrichment was collected to be used for total proteome profiling. Peptides (whole proteome) were fractionated into 24 fractions using basic reverse phase HPLC. 12 fractions were solubilized, desalted by stage tip, and analyzed on an Orbitrap Lumos mass spectrometer. Phosphopeptides were solubilized, desalted by stage tip and analyzed on an Orbitrap Eclipse mass spectrometer with FAIMS enabled. Phosphopeptides were shot twice using different FAIMS CVs. **Data analysis:** MS2 spectra were searched using the COMET algorithm against a Human Uniprot composite database containing its reversed complement and known contaminants. For phospho-proteome, database with phosphorylation on Ser, Thr, or Tyr as differential modifications were searched. For the total proteome, Peptide spectral matches were filtered to a 1% false discovery rate (FDR) using the target-decoy strategy combined with linear discriminant analysis. The proteins were filtered to a <1% FDR and quantified only from peptides with a summed SN threshold of >180. Phosphopeptide FDR was filtered to 1% by LDA analysis of modified peptides only. Phosphosite localization was scored by ModScore. 18-plex Comet Search Parameters: Peptide Mass Tolerance: 50 ppm, Fragment Ion Tolerance: 0.4 ppm (0.02ppm for phosphopeptides), Max Internal Cleavage Site: 2 (3 for phosphopeptides), Max differential/Sites: 5. Methionine oxidation and Phosphorylation at STY are used as variable modification. Localized sites have A-Score >13 with 95% confidence of detection. 18-plex TMTpro Reporter Quant Parameter: Proteome (MS3) - tolerance = 0.003, ms2_isolation_width = 0.7, ms3_isolation_width = 1.2. Phos (MS2) - tolerance = 0.003, ms2_isolation_width = 0.5. Proteins quantified by >1 peptide were analyzed.

### RNA sequencing

RNA library preparation, sequencing and data analysis was conducted at Azenta Life Sciences (South Plainfield, NJ, USA) as follows: **Library Preparation with PolyA selection and Illumina Sequencing:** RNA samples were quantified using Qubit 2.0 Fluorometer (ThermoFisher Scientific, Waltham, MA, USA) and RNA integrity was checked with 4200 TapeStation (Agilent Technologies, Palo Alto, CA, USA). Strand-specific RNA sequencing library was prepared by using NEBNext Ultra II Directional RNA Library Prep Kit for Illumina following manufacturer’s instructions (NEB, Ipswich, MA, USA). Briefly, the enriched RNAs were fragmented for 8 minutes at 94 °C. First strand and second strand cDNA were subsequently synthesized. The second strand of cDNA was marked by incorporating dUTP during the synthesis. cDNA fragments were adenylated at 3’ends, and indexed adapter was ligated to cDNA fragments. Limited cycle PCR was used for library enrichment. The incorporated dUTP in second strand cDNA quenched the amplification of second strand, which helped to preserve the strand specificity. The sequencing library was validated on the Agilent TapeStation (Agilent Technologies, Palo Alto, CA, USA), and quantified by using Qubit 2.0 Fluorometer (ThermoFisher Scientific, Waltham, MA, USA) as well as by quantitative PCR (KAPA Biosystems, Wilmington, MA, USA). The sequencing libraries were multiplexed and clustered onto a flowcell on the Illumina NovaSeq instrument according to manufacturer’s instructions. The samples were sequenced using a 2x150bp Paired End (PE) configuration. Image analysis and base calling were conducted by the NovaSeq Control Software (NCS). Raw sequence data (.bcl files) generated from Illumina NovaSeq was converted into fastq files and de-multiplexed using Illumina bcl2fastq 2.20 software. One mismatch was allowed for index sequence identification. **Data Analysis:** After investigating the quality of the raw data, sequence reads were trimmed to remove possible adapter sequences and nucleotides with poor quality. The trimmed reads were mapped to the reference genome available on ENSEMBL using the STAR aligner v.2.5.2b. The STAR aligner is a splice aligner that detects splice junctions and incorporates them to help align the entire read sequences. BAM files were generated as a result of this step. Unique gene hit counts were calculated by using feature Counts from the Subread package v.1.5.2. Only unique reads that fell within exon regions were counted and reads for each gene were normalized to arrive at a “Transcripts Per Million” (TPM) metric of abundance. Fold change was calculated as (average of the RNA abundance across biological replicates of cells treated with ‘Anti+Oligo’ + 0.1) / (average of the RNA abundance across biological replicates of cells treated with DMSO for 3 days + 0.1).

### Western blot

Equal number of cells from each condition (1-2 x10^6^) were harvested by pelleting at 650 g for 2 min, washed once with PBS, and lysed directly in 150 μL LDS sample buffer (Nupage NP0007) diluted to 2x with water or 100 μl lysis buffer containing 25 mM HEPES, pH7.5, 25 mM KCl, 0.5% NONIDET P40, 1% (v/v) protease inhibitor cocktail (Sigma Cat#P8340) and 1 mM PMSF. Ten percent of the sample (or 40-80 μg protein) was run on Novex™ 4-20% Tris-Glycine Mini Gels (Thermo Fisher Scientific XP04200BOX) or polyacrylamide gels (15% for Tom20, 12% for Tubulin and MMADHC, 6% for MTR) and transferred to a 0.2 μm PVDF membrane (BioRad 1704156). Then, the gel was stained with Coomassie and membrane with Ponceau at the end of the western blot, where shown, to stain total proteins. Membranes were blocked for 10 minutes in 5% non-fat dry milk prepared in TBST. Membranes were then incubated with primary antibody diluted (1:5000 for MMADHC polyclonal, 1:1000 for MMADHC monoclonal, 1:10,000 tubulin and TOM20, 1:1000 for MTR) in 5% milk overnight at 4 °C. The following primary antibodies were used:

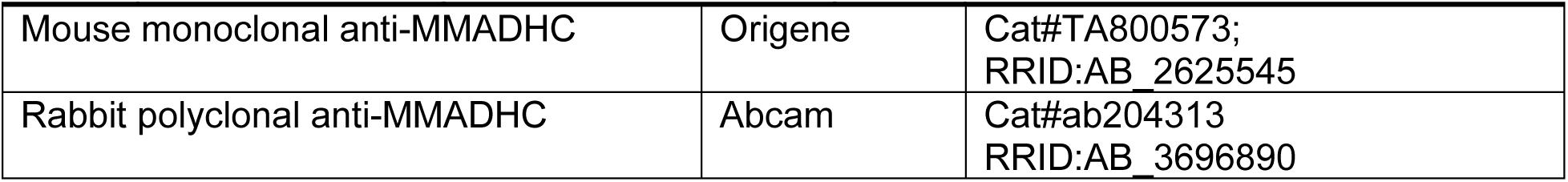
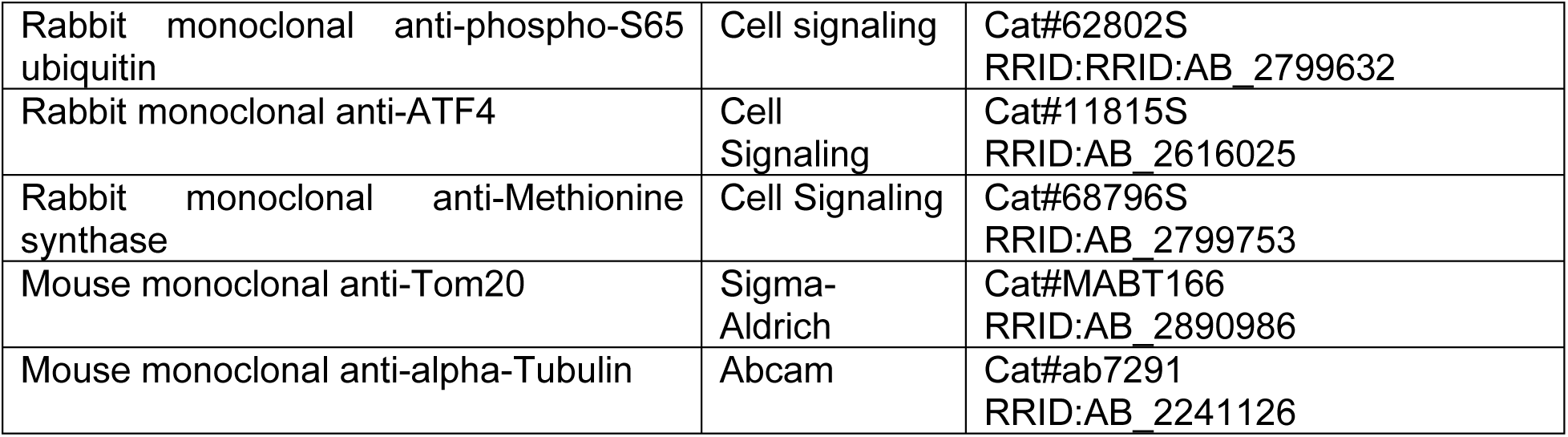

|  |  |  |
| --- | --- | --- |
| Mouse monoclonal anti-MMADHC | Origene | Cat#TA800573;<br>RRID:AB_2625545 |
| Rabbit polyclonal anti-MMADHC | Abcam | Cat#ab204313<br>RRID:AB_3696890 |
| Rabbit monoclonal anti-phospho-S65 ubiquitin | Cell signaling | Cat#62802S<br>RRID:RRID:AB_2799632 |
| Rabbit monoclonal anti-ATF4 | Cell Signaling | Cat#11815S<br>RRID:AB_2616025 |
| Rabbit monoclonal anti-Methionine synthase | Cell Signaling | Cat#68796S<br>RRID:AB_2799753 |
| Mouse monoclonal anti-Tom20 | Sigma-Aldrich | Cat#MABT166<br>RRID:AB_2890986 |
| Mouse monoclonal anti-alpha-Tubulin | Abcam | Cat#ab7291<br>RRID:AB_2241126 |

The primary antibodies used in this work are listed in the Key Resources Table. Membranes were then washed in TBST 4 times for 5 minutes each at room temperature. Next, they were incubated in HRP-linked α-rabbit (Cell Signaling 7074P2 1:5000 or Abcam, ab205719, 1:20,000) or α-mouse (Cytiva NA931V 1:5000, or Kindle Biosciences, R1005, 1:1000) secondary antibodies diluted 1:5000 in 5% non-fat dry milk, for 1 hour at room temperature. Membranes were washed again in TBST as before, incubated with 2 mL of Lightning® Plus ECL (Revvity NEL103E001EA) or Kwikquant Western Blot Detection kit, Kindle Biosciences (Cat#R1004), and exposed to Amhersham Hyperfilm™ (GE Healthcare) or and images were collected with a KwikQuant Imager (Kindle Biosciences). Band intensities were quantified using GelQuant (BiochemLabSolutions) or ImageJ and normalized to total protein.

### Quantitative PCR

RNA was purified using the RNeasy kit (QIAGEN) with on-column DNase (Qiagen ID. 79254) treatment per manufacturer’s instructions. RNA was quantified using a nanodrop and 1 μg was converted to cDNA using the High-Capacity RNA-to-cDNA™ Kit (Applied Biosystems Cat#4387406). QPCR was performed using the primers listed in the Key Resources Table and iQ™ SYBR® Green Supermix (Bio-rad). Fold change was calculated using the ΔΔCt method, normalizing genes of interest to ACTB. Primers used: HSPD1_F: ATGATGCTATGGCTGGAGATT, R: CACCAGCAGCATCCAATAAAG ACTB_F: CACAGAGCCTCGCCTTT, R: GCGGCGATATCATCATCCAT

### Cell fractionation

0.8-1 x10^8^ confluent HEK293 or HT29 cells were washed with PBS and pelleted at 700 g for 5 min. The cells were homogenized in the homogenization buffer containing 20 mM HEPES, pH7.6, 220 mM mannitol, 70 mM sucrose, 1 mM EDTA + 0.5 mM PMSF. The cell debris was removed by centrifugation at 1,000 g for 7 min at 4°C. The crude mitochondria were separated from the cell lysates by the centrifugation at 10,000 g for 15 min and washed once with the homogenization buffer. The isolated cytoplasmic and mitochondrial fractions were frozen in liquid nitrogen and stored at -80°C until further use.

### Target guide cloning

CRISPR guide sequences used were:

MMADHC sgRNA #1 TAACTAAAGAGCAAAATCCT MMADHC sgRNA #2 TGAAAGACATGAGTTTGTGA MTR sgRNA #1 CACCTGCATTGGGATAACAG MTR sgRNA #2 GTGTACGGTCCAATCCTGAA sgOR7G3 sgRNA GGTGAAACAGATGTCGACCA

Guides were cloned according to the protocol available on addgene with pLentiCRISPRv2. Briefly, 5 μg of the vector was digested with FastDigest BsmBI (Fermentas) and dephosphorylated with FastAP (Fermentas) in the FastDigest buffer for 30 min at 37°C. The larger fragment was gel purified using the QIAquick Gel Extraction Kit. Oligos for the guides listed in the key resources table were phosphorylated with T4 PNK for 30 min at 37°C and annealed by heating to 95°C 5 min and then ramping down to 25°C at 5°C/min. The phosphorylated and annealed oligos were diluted 1:200 and ligated to the BsmBI-digested plasmid with Quick Ligase (NEB M2200S) at room temperature for 10 min. The ligation was transformed into Stbl3 bacteria and confirmed by Sanger sequencing.

### Lentivirus production

1 x10^6^ HEK293T cells were seeded in 2 ml media/well in a 6-well plate (three wells per lentivirus). The next day, each well was transfected with 500 μL of transfection mixture. This mixture contained 6 μL Lipofectamine 2000 (Thermo Fisher Scientific Cat#11668019), 1 μg of the lentiviral vector of interest, 900 ng psPAX2 (Addgene 12260), and 100 ng pMD2.G (Addgene 12259) all diluted in 500 uL Opti-MEM medium (Gibco Cat#31985062) and incubated at room temperature for 20 min before adding to cells. Two days after transfection, media was harvested and spun at 650g for 5 min to pellet and eliminate floating cells. The media containing lentivirus was either aliquoted and stored at –80°C or directly used for lentiviral infection.

### Knockout generation

Lentivirus was added with 8 μg/mL polybrene to 2 million K562 (or 1 million HEK293) cells in 2 mL media in one well of a 6-well plate. The plate was spun at 500g for 30 mins (or 1200 g for 45 mins) at room temperature. The next day, infected cells were expanded to a larger culture volume and selected with 2 μg/mL puromycin on day 2-3. KO was confirmed by western blot after treating cells with or without 100 nM antimycin and 30 nM oligomycin, which allows detection of MMADHC in control cells.

### Stable over-expression

The following primers were used to generate the delN38 MMADHC construct from the C2ORF25/MMADHC (NM_015702) Human Tagged ORF Clone (Origene RC204801) F: ATCTGCCGCCGCGATCGCCATGTCGGATGAGTCTCATGTG R: ATCGCGGCGGCAGAT 0.5 x10^6^ HEK293 cells were seeded in 2 mL media/well in a 6-well plate (three wells per lentivirus). The next day, each well was transfected with 250 μL of the transfection mixture in 2 mL media. This mixture contained 3.75 μL Lipofectamine 3000 (Invitrogen Cat#L3000008), 2.5 μg of the vector of interest, 5 μL P3000, all diluted in 250 uL Opti-MEM medium (Gibco Cat#31985062) and incubated at room temperature for 20 min before adding to cells. Two days after transfection, the transfected cells were expanded to a larger culture volume and selected with 100 μg/mL Geneticin (Gibco, Cat#10131035).

### MTR activity assay in cell lysates

For preparing cell lysate, ∼500 µg of cell pellet was resuspended in 1mL of lysis buffer (10 mM Tris-HCl pH - 7.4, 100 mM NaCl, 2 mM ethylenediaminetetraacetic acid (EDTA), 1% Nonidet P-40 (NP-40), 1mM dithiothreitol (DTT) and 1 mM phenylmethylsulfonyl fluoride (PMSF)). 25 µM methylcobalamin (MeCbl) was added to the cell lysate and incubated in the dark on ice for 10 mins, with gentle mixing every 3 mins. The cell lysate was centrifuged at 13000 rpm for 15 mins to pellet cell debris and the supernatant was used to measure MTR activity. MTR activity was measured using the non-radioactive assay as described previously (1). The reaction mixture (800 µL total volume) containing 1mM D,L-homocysteine, 150 µM S-adenosylmethionine (AdoMet), 1.2 µM methionine synthase reductase (MTRR) and 250 µL cell lysate (equivalent to 3 mg of total protein based on Bradford based quantification) in 100 mM potassium phosphate buffer, pH 7.4 was incubated at 37 °C for 5 min following which 300 µM 5-methyltetrahydrofolate (*(6S)*-CH3-H4F) was added and incubated at 37 °C for an additional 2 min. The reaction was initiated by addition of 200 µM NADPH. Aliquots (150 µL) of the reaction mixture were removed at the desired times (typically 5, 10, 15, 20 and 25 min) and quenched by adding 37.5 µL of a 1:1 mixture of formic acid:HCl. The reaction was heated at 80 °C for 10 min and cooled, which led to methenyl-H4F formation. The reaction was centrifuged at 13000 rpm for 7 min and the supernatant was used to determine the concentration of methenyl-H4F at 350 nm using an extinction coefficient of 26.5 × 10^3^ M^−1^ cm^−1^. For each reaction, the initial velocity (V) was obtained from the slope of the line fitted to the product formed versus time plot using the equation:

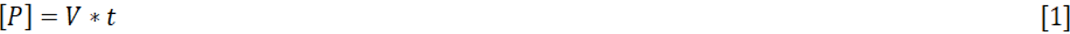

where [P] is the concentration of product formed, and t is the time of the reaction. Deuterated formate tracing K562 cells were treated with DMSO or 100 nM Antimycin + 30 nM Oligomycin for three days and grown in DMEM (Gibco 11995-065) with 10% FBS (Sigma-Aldrich Cat#F2442), 100 U/mL penicillin/streptomycin, and 50 μg/mL uridine. Nine hours prior to harvesting for metabolomics, 1 million cells were washed with, and resuspended in, 2 mL DMEM with 10% dialyzed FBS (Gibco Cat#26400044), 100 U/mL penicillin/streptomycin, 50 μg/mL uridine. Each sample was first split into two groups, receiving 3 mM Sodium formate that was either fully deuterated (Cambridge Isotope Laboratories, Cat#DLM-1361-5) or un-deuterated (Sigma-Aldrich, Cat#247596-100G). Each formate group was further split into two groups: one received supplemental 100 μM homocystine (Sigma-Aldrich H6010) + 100 μM cyanocobalamine (Sigma-Aldrich V6629) and another with only supplemental 100 μM homocystine but no B12 for the duration of labeling (9 hours). To harvest, cells were pelleted at 650g for 2 mins, washed with 1 mL PBS, and spun for 650g 2 mins. PBS was aspirated as close to the cell pellet as possible and tubes were placed on ice immediately. 400 μL of pre-chilled 4:4:2 methanol/acetonitrile/water+0.1M formic acid was added to pellets, tubes were vortexed for 10 seconds, & kept on ice for 3 mins. Samples were neutralized with 35 μL 15% ammonium bicarbonate, vortexed, and stored at –80°C for mass spectrometry analysis. 5 μL of extracted samples analyzed using liquid chromatography coupled with a mass spectrometer (LC-MS). The LC separation was performed on a Waters XBridge BEH Amide column (2.1 mm × 150 mm × 2.5 µm) with column temperature at 25°C. Buffer A was 5% acetonitrile with 20 mM ammonium acetate, 0.25% ammonium hydroxide, pH 9.0 and buffer B was 100% acetonitrile. The total flow rate was set to 0.15 mL/min and the samples were loaded at 90% B. The gradient was, 0 min, 90% B; 2 min, 90% B; 3 min, 75%; 7 min, 75% B; 8 min, 70%, 9 min, 70% B; 10 min, 50% B; 12 min, 50% B; 13 min, 25% B; 14 min, 25% B; 16 min, 0% B, 20.5 min, 0% B; 21 min, 90% B; 25 min, 90% B. The total running time was 25 min. The elute was directed into Q-Exactive Plus Orbitrap mass spectrometer (Thermo Fisher Scientific, Waltham, MA) with HESI probe operating in polarity switching mode. MS parameters were: sheath gas flow 50, aux gas flow 10, sweep gas flow 2, spray voltage 3.3 kV in both negative and positive modes, capillary temperature 310°C, S-lens RF level 50 and aux gas heater temperature 370°C. Other MS parameters included: resolution of 140,000 at m/z 200, automatic gain control (AGC) target at 3e6, maximum injection time of 200 ms and scan range of m/z 120-1000 in the positive mode and m/z 70-1000 in the negative mode respectively. Data were acquired using Xcalibur (v.4.1.31.9, Thermo Fisher). All LC-MS data were collected with samples injected in a randomized order with pooled quality control samples interspersed throughout the runs to quantify instrument performance during the run. Data was analyzed via MAVEN software using an in-house library^38,39^ with 5 ppm mass tolerance and isotope labeling data were corrected for natural isotope abundance^40^.

### Data and code availability

RNA sequencing and TMT proteomics data will be submitted to GEO and ProteomeXchange respectively and be publicly available upon publication. The processed data are also included with manuscript as supplemental information. This paper does not report original code.

## Acknowledgments

We thank all members of the Mootha and Banerjee laboratories for helpful discussions and feedback. We thank Steve Gygi and the TCMP facility at Harvard Medical School for TMT proteomics and Azenta for RNAseq. This work was supported by NIH grants K00CA212468 and K99GM145848 (to S.P.R.) and DK45776 (to R.B.). V.K.M. is an investigator of the Howard Hughes Medical Institute.

## Author Contributions

S.P.R., Z.L., A.G., F.D. designed and performed experiments, S.P.R., Z.L., A.G., F.D., R.B. and V.K.M analyzed data, S.P.R. R.B. and V.K.M conceptualized the study and wrote the original draft, all authors edited the manuscript, R.B. and V.K.M supervised the study.

---

**Fig. S1.**
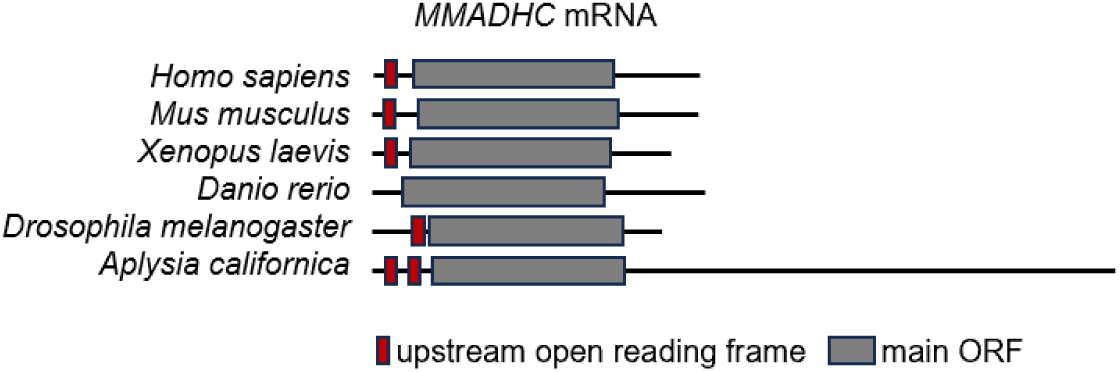
Upstream open reading frames in MMADHC mRNA, related to. **Figure 1**. Schematic of upstream open reading frames (uORFs) in *MMADHC* mRNA in the indicated species. The mRNA and main ORF lengths are drawn to scale, red boxes indicate presence of uORF.

**Fig. S2.**
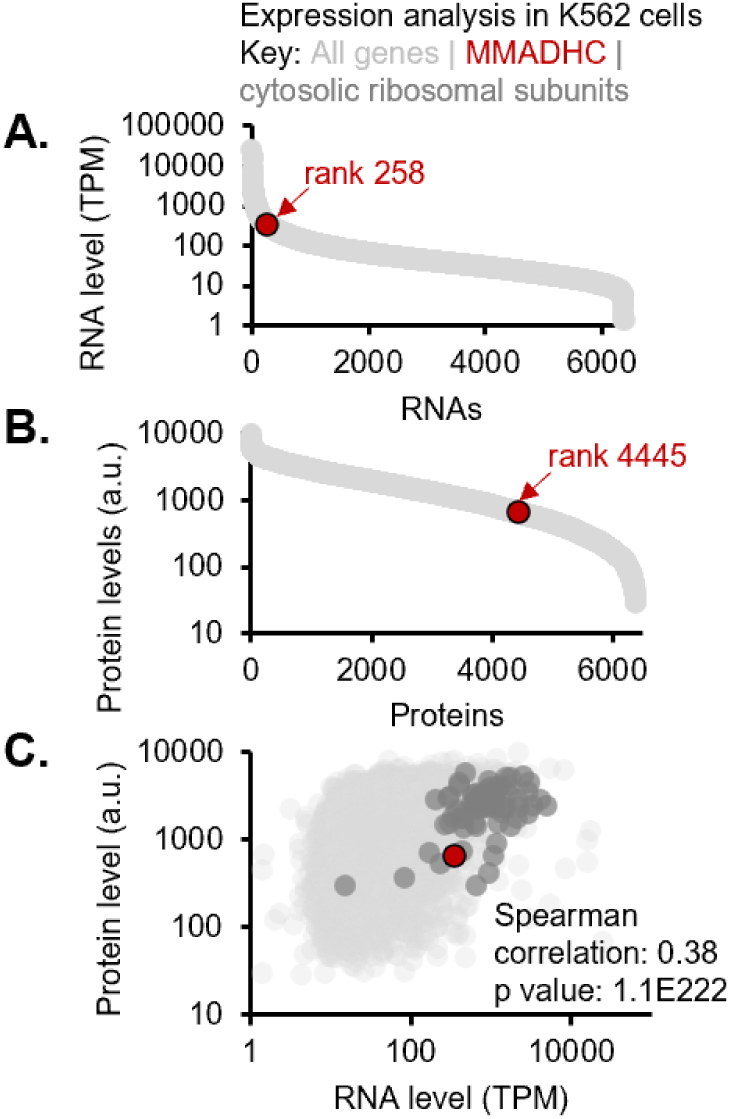
MMADHC protein, but not RNA, is lowly expressed, related to. **Figure 2**. (a) Genes ranked by RNA level in naïve K562 cells measured by RNAseq. (b) Genes ranked by protein level in naïve K562 cells measured by TMT proteomics. Bottom: Scatter plot of protein vs RNA level. Red point corresponds to MMADHC.

**Fig. S3.**
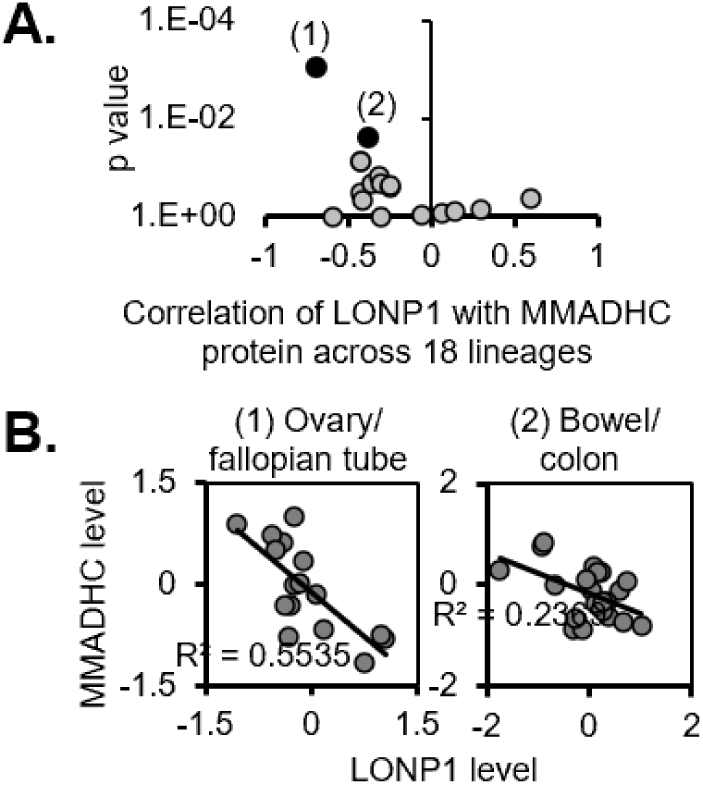
Correlation of MMADHC and LONP1 across CCLE by lineage, related to. **Figure 2**. (A) Volcano plot of correlation between MMADHC and LONP1 in each CCLE lineage. (B) Scatter plots of MMADHC vs LONP1 protein levels in the top two lineages in which the two proteins are most significantly negatively correlated.

**Fig. S4.**
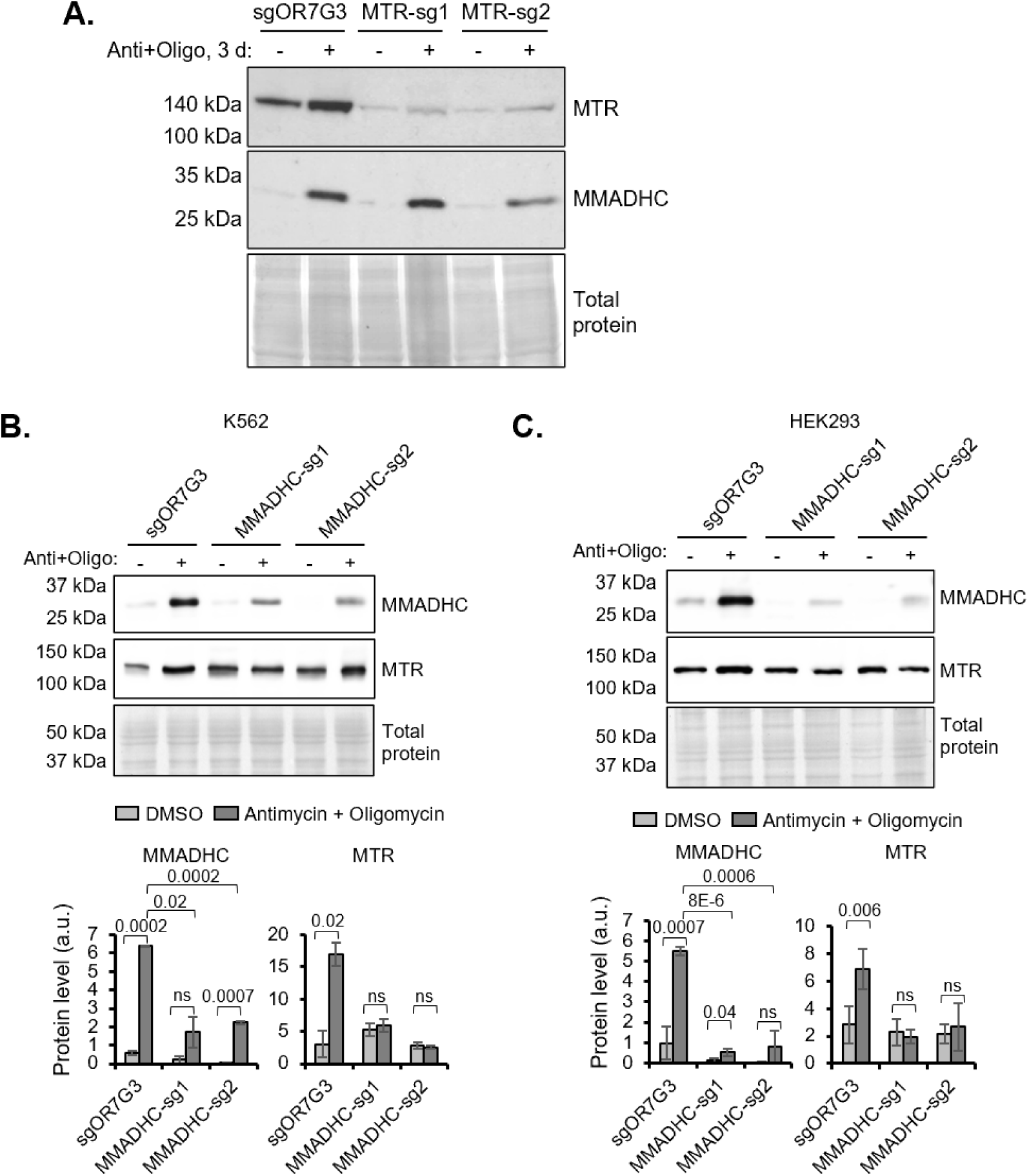
Increase of MTR upon mitochondrial depolarization is MMADHC-dependent and not vice versa, related to. **Figure 3**. (A) Western blot analysis of MMADHC and MTR in cutting control (sgOR7G3) and MTR KO K562 cells treated with antimycin and oligomycin for 3 days. (B, C) Western blot analysis of MMADHC and MTR in cutting control (sgOR7G3) and MMADHC KO K562 (B) and HEK293 (C) cells treated with antimycin and oligomycin for 3 days. Bar plots show quantification of western blots normalized to total protein. Error bars show standard deviation across two to four biological replicates.

**Fig. S5.**
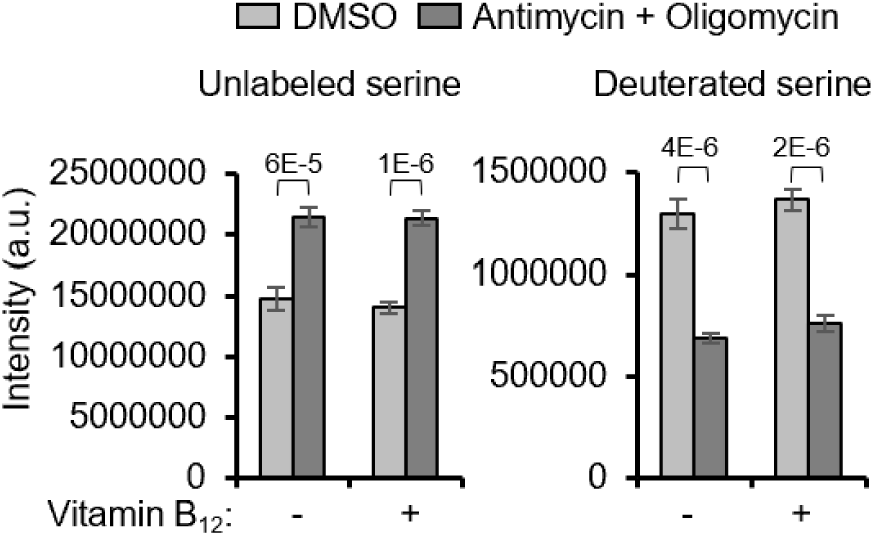
Formate-to-serine flux decreases in cells treated with antimycin and oligomycin, related to Figure 3. Levels of undeuterated and deuterated serine quantified by mass spectrometry in K562 cells treated with antimycin and oligomycin for 3 days and labelled with fully deuterated formate for 9 hours in the presence or absence of vitamin B12. Error bars represent standard deviation of four biological replicates.

